# Imaging *Clostridioides difficile* spore germination and germination proteins

**DOI:** 10.1101/2022.05.31.494260

**Authors:** Marko Baloh, Hailee N. Nerber, Joseph A. Sorg

**Author notes:** Corresponding author, Em.

## Abstract

*Clostridioides difficile* spores are the infective form for this endospore-forming organism. The vegetative cells are intolerant to oxygen and poor competitors with a healthy gut microbiota. Therefore, in order for *C. difficile* to establish infection, the spores have to germinate in an environment that supports vegetative growth. To initiate germination, *C. difficile* uses Csp-type germinant receptors that consist of the CspC and CspA pseudoproteases as the bile acid and co-germinant receptors, respectively. CspB is a subtilisin-like protease that cleaves the inhibitory pro-peptide from the pro-SleC cortex lytic enzyme thereby activating it and initiating cortex degradation. Though several locations have been proposed for where these proteins reside within the spore (*i*.*e*., spore coat, outer spore membrane, cortex, inner spore membrane), these have been based, mostly, on hypotheses or prior data in *C. perfringens*. In this study, we visualize the germination process using TEM and SEM, and, using immunogold labeling of the spore proteins, find that these proteins are localized to the spore cortex, consistent with the observed, rapid, changes to the spore structure during germination.

**Importance:** Germination by *C. difficile* spores is the first step in the establishment of potentially life-threatening CDI. A deeper understanding of the mechanism by which spores germinate may provide insight for how to either prevent spore germination into a disease-causing vegetative form, or trigger germination prematurely when the spore is either in the outside environment or in a host environment that is non-conducive to the establishment of colonization / disease.

## Introduction

*Clostridioides difficile* is a Gram-positive, obligately anaerobic, spore-forming bacterium that is opportunistically pathogenic in humans. *C. difficile* vegetative cells are susceptible to oxygen and therefore incapable of surviving outside of the host gastrointestinal (GI) tract (1, 2). The spore form is the metabolically dormant stage in the *C. difficile* lifecycle and persists outside of the host in the oxygen-rich environment. Disruptions to the normally-protective native gut microbiota are commonly the result of broad-spectrum antibiotic use (3-6). While the *C. difficile* vegetative form is susceptible to antibiotics, the spores are not and therefore may remain in the gut to initiate recurring infection (7).

*C. difficile* spores, like spores from other spore-forming organisms, have a complex structure that conveys resistance to various environmental conditions inside and outside of the body (8, 9). The spore core contains the DNA, RNA, and protein required for resumption of metabolic processes upon germination. It is also rich in pyridine-2,6-dicarboxylic acid (dipicolinic acid [DPA]), chelated with calcium (Ca-DPA). DPA is packaged into the core during sporulation in exchange for water, and the high DPA / low water content of the core conveys the dormant spores with remarkable heat resistance (9-17). The core is surrounded by the inner spore membrane with low permeability to water. The inner spore membrane is surrounded by the germ cell wall, a layer of peptidoglycan that becomes the cell wall of a newly germinated cell (18, 19). Surrounding this is the cortex, a thick layer of modified peptidoglycan where much of the muramic acid residues are converted to the muramic-δ-lactam moieties that are the target of the cortex lytic enzymes (20-24). Surrounding the cortex is the outer spore membrane (25, 26), the coat layer (27), and the exosporium (28-33).

Upon entry into a susceptible host, *C. difficile* spores germinate in response to small molecule germinants (34). *C. difficile* spores respond to host-derived bile acids, synthesized by the liver using cholesterol as a scaffold, as activators or inhibitors of germination (35, 36). In the liver, the primary bile acids cholate and chenodeoxycholate are modified by conjugation with either taurine or glycine (37, 38). Cholate derivatives, taurocholate in particular, are the most effective *C. difficile* spore germinants, but chenodeoxycholate is a competitive inhibitor of cholic acid-mediated germination (35, 36, 39-42). Though these primary bile acids are required, they are not sufficient to trigger *C. difficile* spore germination. Amino acids and calcium can function as co-germinants, and glycine has been shown to be the most efficient co-germinant, especially *in vivo*, since the depletion of glycine in mouse models prevents the establishment of CDI (35, 41, 43-45).

In the majority of the studied endospore-forming organisms, spores detect germinants via Ger-type receptors embedded in the inner spore membrane (46). *C. difficile* does not encode Ger-type germinant receptors and, instead, uses the CspA and CspC proteins as germinant receptors (47-51). Prior to the work done in *C. difficile*, CspA, CspB, and CspC proteins were characterized in *Clostridium perfringens*, and found to be subtilisin-like serine proteases. These proteins contain a catalytic triad that is capable of processing the cortex lytic enzyme SleC from its inactive pro-form into an active enzyme that degrades the cortex and initiates spore germination (52, 53). Unlike what is found in *C. perfringens, C. difficile cspB* and *cspA* are translationally fused (CspBA), and *cspC* is encoded downstream from the *cspBA* gene. Interestingly, both CspA and CspC have lost their catalytic triad and, therefore, are pseudoproteases (47, 48).

Translation of *cspBA* produces CspBA which then undergoes interdomain processing by YabG to generate the CspB protease and the CspA pseudoprotease. When activated, CspB cleaves the inhibitory pro-peptide from pro-SleC to trigger germination (49-51). The disruption of *cspA* prevents incorporation of CspC into the developing spore, suggesting interaction of CspA and CspC and cleavage of CspA from CspBA is important for response to co-germinants, suggesting that CspA functions as the co-germinant receptor (49, 50, 54). Finally, CspC has been identified to be the bile acid germinant receptor or as a hub for germinant processing (36, 39, 48, 55).

There are several hypotheses about where the Csp-type germinant receptors are located in *C. difficile* spores. Prior studies have proposed locations in the spore coat and outer spore membrane (56), outer spore membrane alone (57), or the inner spore membrane (9), where they exist as either a complex or individually. Here, we used TEM imaging and immunogold labeling to show that CspB, CspA, CspC, and SleC are localized to the spore cortex layer. This is consistent with our observation that the cortex thickness decreases dramatically within 5 minutes of the initiation of germination. We also show by SEM that germination and the transition of a spore into a vegetative cell can proceed entirely while the nascent cell is still enveloped in the coat and exosporium layer, without observable disruption to those layers.

## Results

### *C. difficile* cortex degradation is a quick event

*C. difficile* germination begins with the detection of germinants and co-germinants by the germinant receptors. This results in the initiation of cortex degradation, release of DPA, core rehydration, and the resumption of metabolic activity. In prior work, we found that DPA release is a quick event, with the majority of DPA being released within 20 minutes of germination initiation (10, 12, 16). Because the DPA release from the spore core is dependent on the activation of the SpoVAC mechanosensing protein embedded in the inner spore membrane (16), the cortex degradation must be a quick event; indeed the cortex fragments are quickly detected after initiation of germination (58). In order to observe the structural changes that occur in the germinating spore (from the early stage of germination to the development of vegetative cell), we induced germination in spores derived from the wild-type *C. difficile* R20291 strain by incubating them with taurocholate (TA) and glycine in BHIS liquid medium. We then sampled between T=0 minutes (ungerminated spores) to T=180 minutes, when we expected to see fully developed vegetative cells, based on prior work (10). The samples were fixed and processed for TEM imaging. In order to observe the potential changes to the morphology of the spore surface, samples were also prepared for SEM imaging.

At T=0 all of the spore structures are clearly distinguishable by TEM (i.e., exosporium, coat, outer spore membrane, cortex, germ cell wall, inner spore membrane, and the core) (Figure 1 A-C). SEM imaging showed no changes in the spore morphology between T=0 and T=60 minutes (Figure 1 D-E). At T=180, we began to observe vegetative cells exiting spores (Figure 1 F). For comparison, we also imaged cultures by both TEM and SEM grown overnight in rich media (BHIS) (Figure 1 G-I).

**Figure 1:**
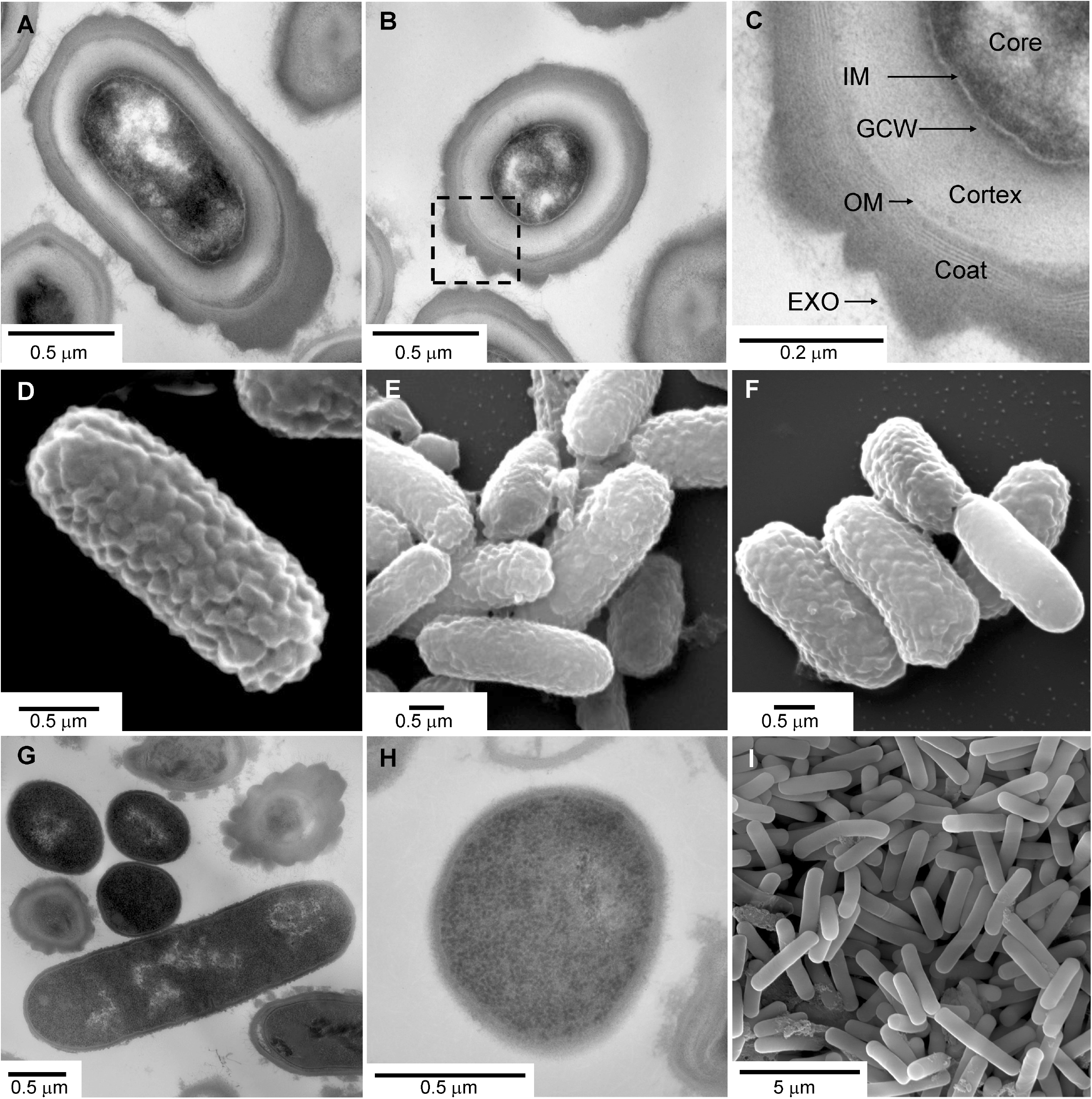
*C. difficile* spore structure and cell morphology visualized by TEM and SEM. Spores (A-F) and vegetative cells (G-I) derived from the *C. difficile* R20291 strain were embedded in epoxy resin, sectioned, and imaged by TEM (A, B, C, G, H) or were chemically dried, sputter coated, and imaged by SEM (D, E, F, I).

Compared to dormant spores (T=0), at T=5, we observed a reduction in the thickness of the cortex, as well as the disappearance of a clearly distinguishable germ cell wall layer (Figure 2 A). Cortex thickness was measured in dormant spores in both horizontally cross-sectioned spores and spores sectioned along the longitudinal axis. These were compared 5 minutes post-germination. The average cortex thickness in both orientations was nearly the same (Figure 2 B). After 5 minutes of incubation in media containing germinants, the average cortex thickness was reduced by approximately 68% for cross-sectioned spores and by 67% for longitudinally sectioned spores (Figure 2 B). These results confirm prior observations by TEM that cortex is rapidly degraded upon germinant sensing.

**Figure 2:**
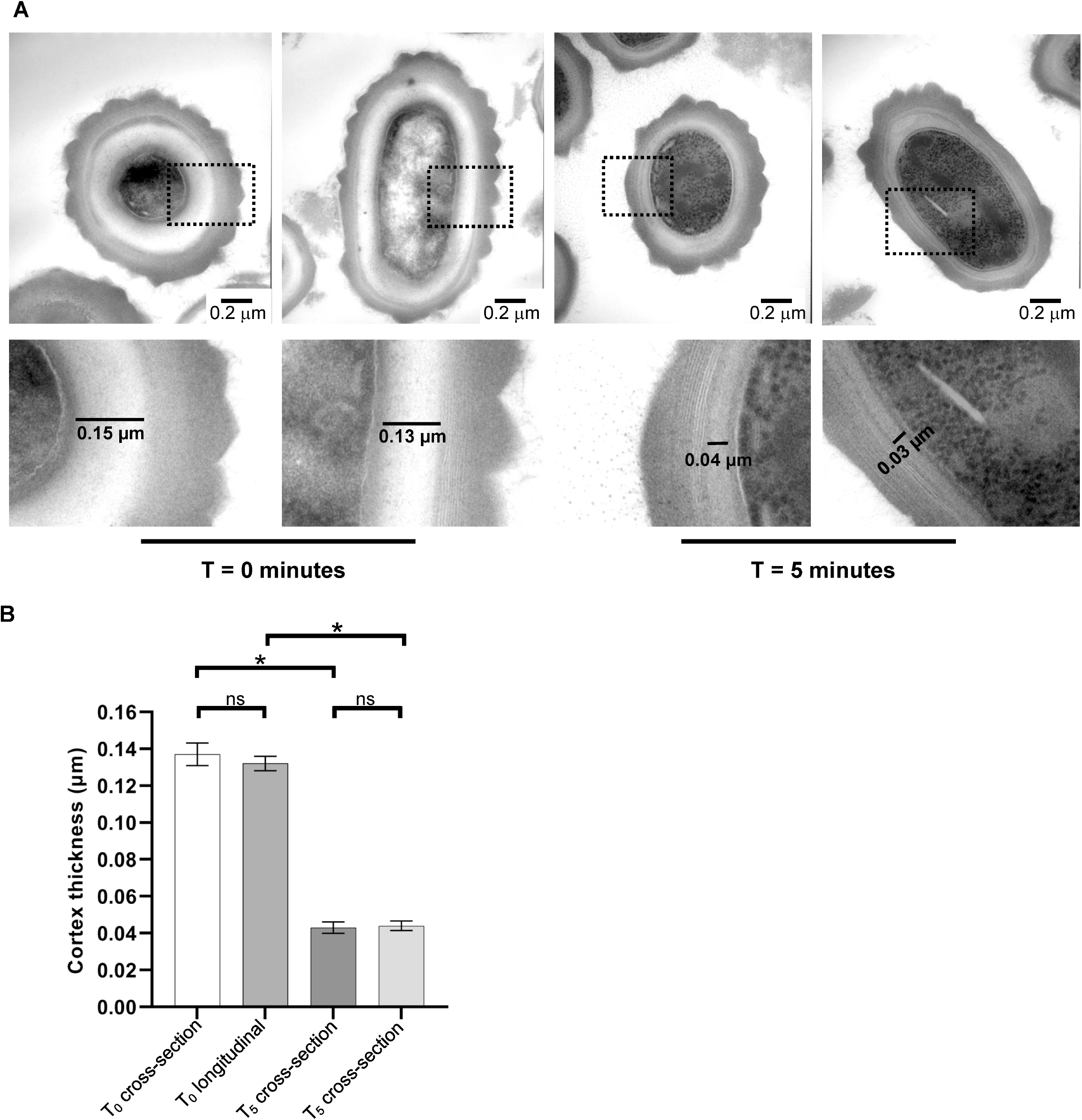
*C. difficile* cortex significantly thins within 5 minutes of germinant addition. *C. difficile* spores were germinated in rich medium supplemented with 10 mM TA and 30 mM glycine. A sample was taken prior to the addition of germinants (T_0_) and 5 minutes post-germinant additions (T_5_). (A) Representative images of T_0_ spores and T_5_ spores. Boxes represent areas of cortex thickness measurement, with magnified view with measurement scale bars shown underneath. (B) Average cortex thickness was measured in cross-sectioned and longitudinally-section spores, at T_0_ and T_5_. N=10 spores of each section orientation counted in both conditions (Total N=20). Cortex thickness was measured using Fiji Scale Bar tool. * indicates p < 0.0001 as determined by one-way ANOVA using Tukey’s multiple-comparison test. ‘ns’ indicates no significance.

Starting at T=5 minutes after germination initiation (Figure 3), the majority of the spores showed reduced cortex thickness, along with unidentified changes within the core. At later time points of germination there is a considerable variation in the spore core appearance. At T=60 minutes we began to observe what looked like early vegetative forms, but still encased inside the spore coat (Figure 3). At T=90 minutes we observed free cells and cells in the process of exiting the coat / exosporium layer (Figure 4). This was more commonly observed event at T=180 (Figure 4). At T=180 minutes, we observed vegetative cells without the coat layer, vegetative cells undergoing division, and spores that had not fully germinated. (Figure 4). This probably reflects the previously established findings that there is considerable variability in spore development (59, 60). Nevertheless, it seems that cortex degradation observed at T=5 occurs in the majority of the spore population.

**Figure 3:**
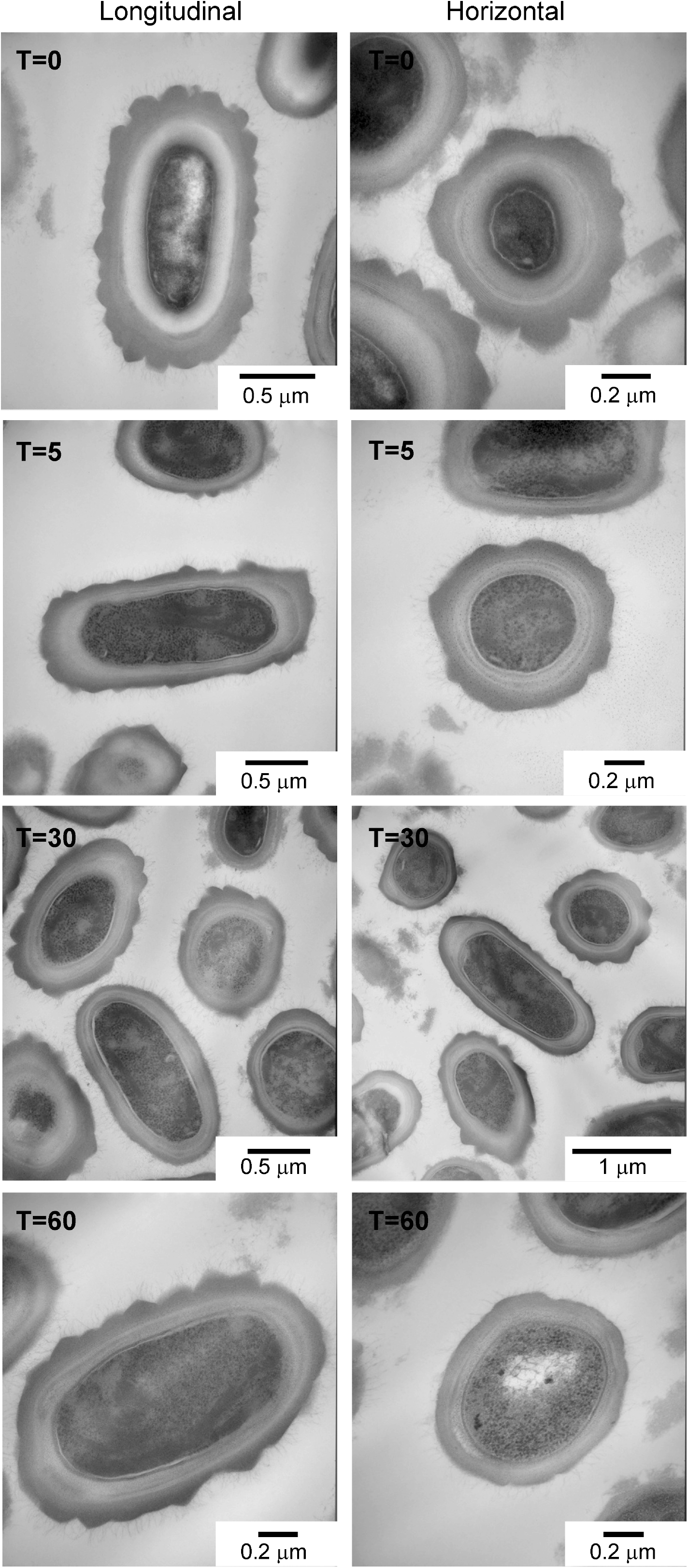
Monitoring *C. difficile* spore germination over time (T_0_ – T_60_). Dormant (T_0_) *C. difficile* R20291 spores, or spores germinated for T_5_, T_30_, and T_60_ minutes were embedded in epoxy resin, sectioned, and imaged by TEM.

**Figure 4:**
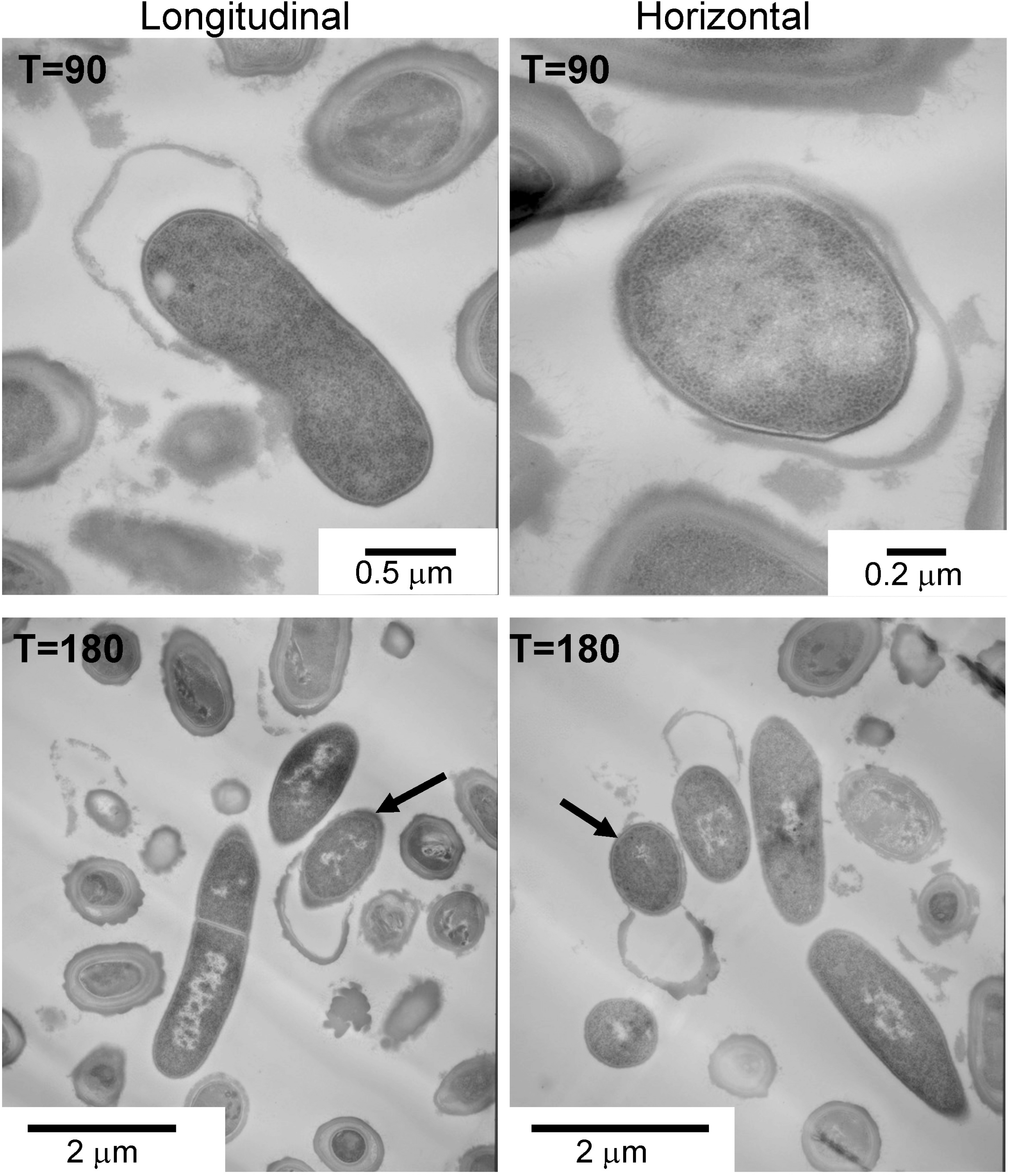
Monitoring *C. difficile* spore germination over time (T_90_ & T_180_). *C. difficile* R20291 spores germinated for T_90_ and T_180_ were embedded in epoxy resin, sectioned, and imaged by TEM. Arrows indicate cells exiting the coat / exosporium layer.

### Germinating *C. difficile* spores develop into vegetative cells inside the protective coat and exosporium layers

While there is no observable difference via SEM imaging between ungerminated spores and spores undergoing germination at the time points tested, T=0, T=5, T=20 (data not shown), we observed cells exiting the exosporium at T=180. We note that the newly hatched vegetative cells exit the exosporium at a pole of the spore, and that the exosporium structure remains intact during the entire germination process (Figure 5). This is not always the case in TEM images, probably due to the different, considerably longer and harsher, sample preparation steps required for TEM that may damage the exosporium layer. Nevertheless, a similar process can be observed in the T=180 spores imaged by TEM (Figure 4).

**Figure 5:**
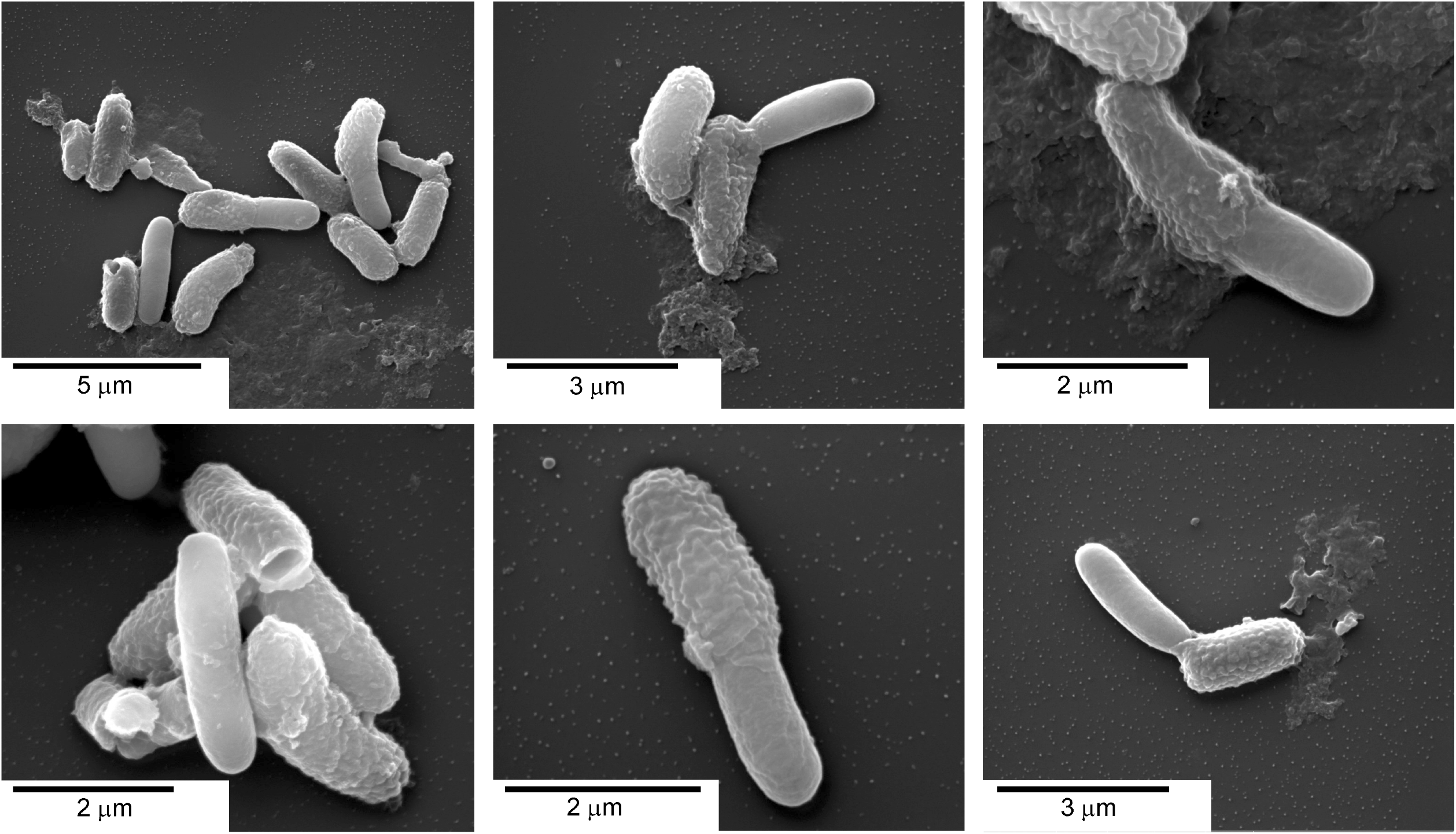
*C. difficile* outgrowth occurs inside the coat / exosporium layers. Representative images of spores and vegetative cells derived from the *C. difficile* R20291 strain, germinated for 180 minutes, fixed, chemically dried, sputter coated, and imaged by SEM.

### CspBA, CspC and SleC are localized to the spore cortex

Our TEM imaging showed that the cortex layer undergoes dramatic thinning within 5 minutes of the initiation of germination. This rapid time frame might suggest the proteins responsible for cortex degradation are already present in this layer. We therefore set out to determine the location of the *C. difficile* CspB, CspA, CspC, and SleC proteins in the spore using immunolabeled TEM samples. To accomplish this, we used two approaches. We embedded spores derived from the wild-type *C. difficile* R20291 and (as negative controls) *C. difficile* R20291 Δ*cspBAC* and *C. difficile* Δ*sleC* in Lowicryl HM20 resin, using high pressure freezing (HPF), then sectioned them. The 70-80 nm sections were labeled with primary antibodies raised against *C. difficile* CspB, CspC, CspA, and SleC, followed by the gold-conjugated secondary antibodies. We also constructed plasmids harboring *cspBA-*FLAG and *sleC*–FLAG tag C-terminal fusions that we transformed into *C. difficile* R20291 Δ*cspBAC* and *C. difficile* Δ*sleC* strains, respectively. These transformed strains were then embedded in resin, sectioned, incubated with primary antibodies targeting the FLAG reporter, followed by secondary gold-conjugated antibodies. In this case, the negative control was *C. difficile* R20291 spores. Samples were imaged, gold labels counted and compared to counts obtained from the negative controls.

Our results show that in both cases, either by direct labeling of the *C. difficile* CspB, CspA, CspC, and SleC proteins, or as in CspBA-FLAG and SleC-FLAG-tagged strains, there is a significant increase in labeling in the spore cortex, compared to the negative controls (Figure 6). Importantly, we did not observe much labeling in the coat or core layers. We conclude that *C. difficile* CspB, CspA, CspC, and SleC are localized to the spore cortex, and are responsible for the observed cortex thinning during the early stages of germination.

**Figure 6:**
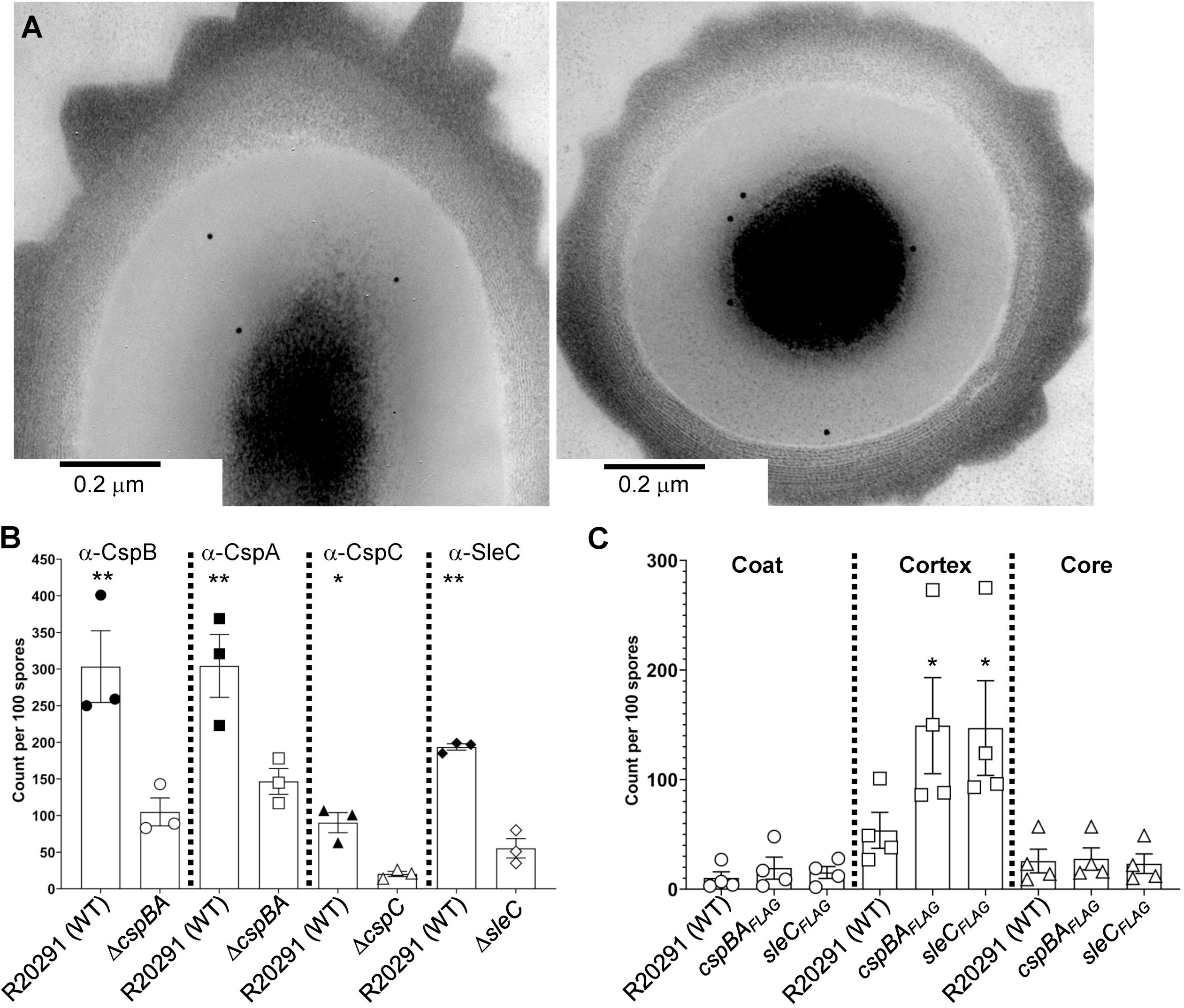
CspB, CspA, CspC, and SleC are located within the spore cortex. Spores derived from *C. difficile* R20291, *C. difficile* Δ*cspBAC, C. difficile* Δ*sleC, C. difficile* Δ*cspBAC* p*cspBA*_*FLAG*_, *and C. difficile* Δ*sleC* p*sleC*_*FLAG*_, were fixed, embedded in acrylic resins, sectioned, immunolabeled with antibodies specific for the CspB, CspA, CspC, SleC proteins or the FLAG epitope and then imaged by TEM. (A) Representative images of immunolabeled spores. (B) Counts for the spores derived from the *C. difficile* R20291 strain, *C. difficile* Δ*cspBAC* and *C. difficile* Δ*sleC* strains labeled with α-CspB, α-CspA, α-CspC, and α-SleC primary and gold-conjugated secondary antibodies. For clarity, only the counts for the particles located in the cortex are shown since the counts for the coat layer and the core do not reach statistical significance. Counts were compared to the counts observed in the deletion mutant. (C) Count for the spores derived from the *C. difficile* R20291, *C. difficile* Δ*cspBAC* p*cspBA*_*FLAG*_, and *C. difficile* Δ*sleC* p*sleC*_*FLAG*_, labeled with α-FLAG primary and gold-conjugated secondary antibodies. The data represent the average of 3 or 4 biological replicates and the error bars represent the standard errors of the mean. **, p < 0.0001, * p < 0 .05 as determined by one-way ANOVA using Šídák’s multiple-comparison test.

## Discussion

*C. difficile* spore germination is a tightly regulated process evolved to ensure that the vegetative cells emerge in an environment that is conducive to survival. For *C. difficile*, the obligately anaerobic cells do not survive exposure to the oxygen-rich environment (along with other potentially damaging conditions like UV radiation, desiccation, etc.). Another challenge is the inability of the *C. difficile* cells to compete against the native host microbiota in healthy individuals, requiring the ability of the metabolically dormant spores to probe for appropriate conditions when ingested by a host. These evolutionary pressures have led to the development of germination pathways that are triggered only when the proper location (the lower gastrointestinal tract of a host organism), and the proper conditions (the lack of competing microbiota) are detected.

*C. difficile* utilizes the Csp-type germinant receptors. There has been considerable research done on the role and the location of *C. perfringens* CspB, CspA, and SleC. Several studies have suggested, using immunoblotting and immunoelectron microscopy, that *C. perfringens* SleC is localized to the spore coat (61, 62). Other studies, again using immunoblotting of the spore coat fraction, have reported this while also suggesting the same location for CspB (52, 53, 63). Though it is plausible that the CspB, CspA, CspC, and SleC proteins are located in the same spore layer in both *C. perfringens* and in *C. difficile*, our results suggest that is not the case, at least for *C. difficile*. The studies that used immunoelectron microscopy to determine the location of SleC have drawn their conclusions from the subjective interpretation of the EM images of the spores labeled with anti-SleC antiserum, without using deletion mutants as negative controls. Here, our approach was to quantify and analyze the labeling differences between the spores derived from *C. difficile* R20291 and the spores derived from the *C. difficile* Δ*cspBAC* and *C. difficile* Δ*slec* strains. Additionally, we constructed CspBA*-*FLAG and SleC-FLAG fusions for further validation and obtained similar results. In the prior work that used immunoblotting of spore coat fractions (52, 53, 61, 63), it is perhaps inevitable that cortex localized proteins are also extracted, which could then lead to the conclusion that the coat is the location of both SleC and CspB. The data from these studies has led to the proposals that Csp proteins and SleC (in its pro-SleC form) are found in the spore coat and / or the spore outer membrane, in a complex or individually (44, 51, 57, 64). Another proposed model states that CspB, CspA, and CspC are in a ‘germinosome’ complex, similarly to what is found in *B. subtilis* (54). In this scenario, the CspB protease activity is inhibited by the CspC bile acid receptor and the CspA co-germinant receptor until the detection of germinants, when it is capable of cleaving the proSleC prodomain, ensuring tight control of germination initiation. SleC is unlikely to be a part of this complex, due to its much greater abundance in the spore (42), but it is reasonable to hypothesize that it is located in the spore cortex, as well. Here, using combined TEM ultrastructure imaging of *C. difficile* spores during the early stages of germination, coupled with TEM immunogold labeling, we show that all three members of the *C. difficile* germinosome, and SleC, indeed are located in the spore cortex.

Our combined TEM / SEM images of germinating *C. difficile* spores have led to an interesting observation that spores fully germinate while still encased in the exosporium layer. This leads to an interesting question of how newly developed cells exit this layer and why it occurs at the spore pole. It has been suggested that the exosporium has a role in adherence of *C. difficile* spores and its experimental removal reduces their adherence to Caco-2 cells, presumably due to the removal of specific proteins that recognize the apical cell surfaces (28). Any other role of the exosporium at this time is unknown, but our finding that it remains undisturbed throughout the germination process *in vitro* tempts us to propose that it may serve a protective role for a newly germinated cell at the earliest stage. We also note that newly hatched cells exiting the exosporium tend to be sickle-shaped, unlike mature vegetative cells (Figure 4), perhaps indicative of the physical stress of exiting the exosporium. Future research into this stage of cell development may provide insight into the mechanisms of exosporium / coat shedding.

## Materials and Methods

### Bacteria and strains

*C. difficile* R20291, and derived strains, were grown at 37 °C in an anaerobic chamber (Coy Laboratories; model B; >4% H2, 5% CO2, 85% N2) on brain heart infusion agar supplemented with 5 g / L yeast extract and 0.1% L-cysteine (BHIS) or TY agar medium (30 g / L Bacto typtone, 20 g / L yeast extract), as indicated. *Escherichia coli* DH5α (65) was grown on LB medium. Chloramphenicol (20 mg / ml), thiamphenicol (10 mg / ml), kanamycin (50 mg / ml), and ampicillin (100 mg / ml) were added where indicated. Spores were generated on 70:30 agar medium (63 g / L Bacto peptone, 3.5 g / L proteose peptone, 11.1 g / L BHI, 1.5 g / L yeast extract, 1.06 g / L tris base and 0.7 g / L ammonium sulfate [(NH_4_)_2_SO_4_].

### Construction of R20291 Δ*cspBAC* and FLAG-tag reporter strains

*C. difficile* R20291 Δ*sleC* strain, a TargeTron *sleC* insertion mutant, was previously constructed by our lab (54). *C. difficile* R20291 Δ*cspBAC* strain, a complete CRISPR-induced deletion of the csp*BAC* operon was constructed, as follows. Plasmid pHN168 was generated by inserting CRISPR_CspB_2 gBlock into PCR 2.1 – TOPO plasmid. pHN168 was digested at the *KpnI* and *MluI* restriction sites to generate a free fragment consisting of the gRNA. This fragment was ligated at 16 °C overnight into pKM126 that had also been digested at the *KpnI* and *MluI* restriction sites. The resulting plasmid was digested with NotI and XhoI to insert the homology arms for the *cspBAC* deletion. The upstream homology fragment was amplified with primers 5’cspBAC_UP and 3’cspBAC_UP while the downstream homology fragment was amplified with 5’cspBAC_DN and 3’cspBAC_DN. Gibson assembly was used to insert these homology fragments into the above digested plasmid, resulting in pHN23. Due to problems with the *tetR* promoter having leaky expression of *cas9*, this was exchanged for the *xylR* promoter. The *xylR* promoter was amplified from pCE641 with the primers 5’cspBAC_HR_xylR and 3’cas9_Pxyl 2. This fragment was inserted into PacI and XhoI digested pHN23 using Gibson assembly (66), resulting in pHN36. HNN08 (R20291 Δ*cspBAC*) was generated by passaging R20291 pHN36 4 times on TY supplemented with Tm and 1% xylose. The resulting strain was cured of the plasmid by passaging twice in BHIS broth (without antibiotic selection).

FLAG-tagged reporter strains were constructed by FLAG-tag fusions to the C-terminal of *C. difficile cspBA* and *sleC* genes on a plasmid that was transformed into the *C. difficile* R20291 Δ*cspBAC* and Δ*sleC* mutant strains, respectively. For CspBA-FLAG fusion, the wild-type *cspBA* genes (including the 500 bp upstream region) were amplified from the *C. difficile* R20291 using primers 5’ CspBA promoter NdeI and 3’ CspBA-FLAG and, along with the gBlock containing FLAG tag, and inserted by Gibson assembly (66) into plasmid pMTL 84151 digested with NdeI / XhoI. For the SleC-FLAG fusion, the wild-type *sleC* gene (including the 500 bp upstream region) was amplified from *C. difficile* R20291 genome using primers 5’ SleC promoter NdeI and 3’ SleC-FLAG, along with with the gBlock containing FLAG tag, and inserted by Gibson assembly into plasmid pMTL 84151 digested with NdeI / XhoI. This yielded plasmids pMB96 and pMB98, respectively. The plasmids were then transformed into *E. coli* DH5α for storage. These plasmids were subsequently transformed into *E. coli* HB101 / pRK24 and grown on LB medium supplemented with chloramphenicol and ampicillin. The resulting strain was grown overnight and then mixed with the *C. difficile* R20291 Δ*cspBAC* or Δ*sleC* strain grown in BHIS in an anaerobic chamber. The conjugation mixtures were spotted onto BHI plates and allowed to grow for 24 hours. Subsequently, the cells were washed with BHIS, and the slurry was transferred onto BHIS medium supplemented with thiamphenicol (for plasmid maintenance) and kanamycin (to counter select *E. coli* growth) [BHIS(TK)]. Thiamphenicol resistant colonies were tested for the presence of the desired fusion construct on the plasmid by PCR and confirmed by sequencing. All strains and plasmids in this study are listed in Table 1. Primers used to construct the strains and plasmids are listed in Table S1 in the supplemental material.

**Table 1.**
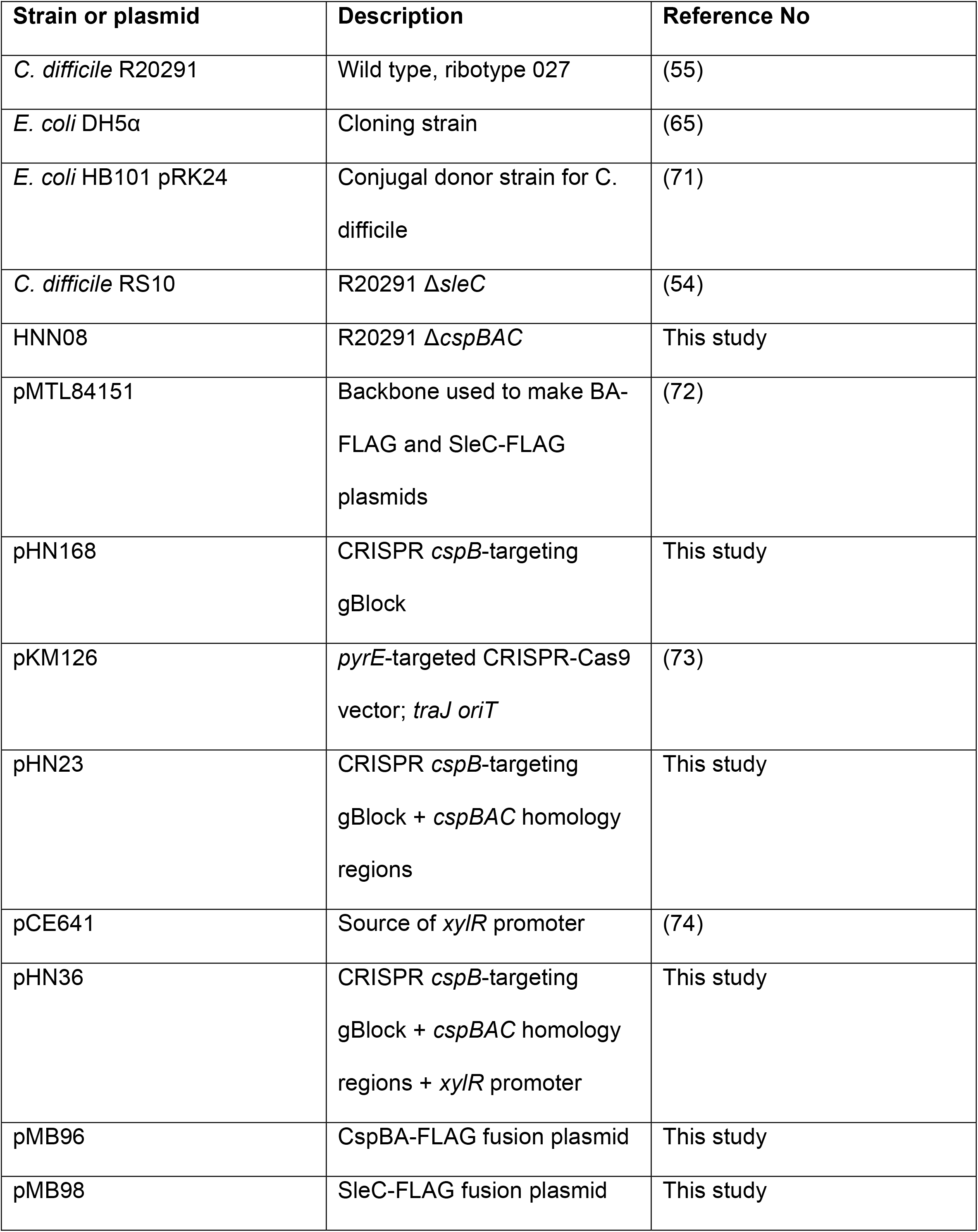
List of strains and plasmids.

| Strain or plasmid | Description | Reference No |
| --- | --- | --- |
| *C. difficile* R20291 | Wild type, ribotype 027 | (55) |
| *E. coli* DH5α | Cloning strain | (65) |
| *E. coli* HB101 pRK24 | Conjugal donor strain for *C. difficile* | (71) |
| *C. difficile* RS10 | R20291 $\Delta sleC$ | (54) |
| HNN08 | R20291 $\Delta cspBAC$ | This study |
| pMTL84151 | Backbone used to make BA-FLAG and SleC-FLAG plasmids | (72) |
| pHN168 | CRISPR *cspB*-targeting gBlock | This study |
| pKM126 | *pyrE*-targeted CRISPR-Cas9 vector; *traJ oriT* | (73) |
| pHN23 | CRISPR *cspB*-targeting gBlock + *cspBAC* homology regions | This study |
| pCE641 | Source of *xylR* promoter | (74) |
| pHN36 | CRISPR *cspB*-targeting gBlock + *cspBAC* homology regions + *xylR* promoter | This study |
| pMB96 | CspBA-FLAG fusion plasmid | This study |
| pMB98 | SleC-FLAG fusion plasmid | This study |

### Spore purification

Spores were purified as previously described (10, 16, 35, 64). Briefly, strains were grown on 70:30 sporulation medium. After 5 days, growth from 2 plates each was scraped into 1 mL distilled water (dH_2_O) in microcentrifuge tubes and left overnight at 4 °C. The cultures were then resuspended in the dH_2_O in the same microcentrifuge tubes and centrifuged at >14,000 × g for 10 min, the top layer containing vegetative cells and cell debris was removed by pipetting, and the rest of the sediment resuspended in fresh dH_2_O. The tubes, again, were centrifuged for 1 min at >14,000 × g, the top layer removed, and the sediment resuspended. This was repeated 5 more times, combining the sediment from 2 tubes into one. The spores were then separated from the cell debris by centrifugation through 50% sucrose solution for 20 min at 4 °C and 3,500 × g. The resulting spore pellet was then washed 5 times with dH_2_O, resuspended in 1 mL dH_2_O, and stored at 4 °C until use.

### TEM ultrastructure sample preparation

For ultrastructure imaging of germinating spores, inside the anaerobic chamber, 50 µL of purified spores was incubated in 950 µL of BHIS supplemented with 10 mM TA and 30 mM glycine inside a 2.0 mL microcentrifuge tube. So that the samples were ready for the subsequent critical fixation step at the same time, the first sample incubated was the latest time point (*i*.*e*., T=180 minutes) and each subsequent tube was a previous time point (*i*.*e*., T=120, T=90, T=60, T=30, T=20, T=10, T=5), with the last tube incubated representing T=0 (ungerminated spores) which contained 950 µL of BHIS without TA or glycine. The tubes were taken out of the anaerobic chamber, centrifuged for 1 min at >14,000 × g, supernatant removed and replaced with 1,950 µL of fixative (5% glutaraldehyde, 2% acrolein in 0.05 M HEPES, pH 7.4). The samples were incubated in the fixative overnight at 4 °C. The following day, the fixed samples were centrifuged for 5 min at >14,000 × g, and post-fixed and stained for 2 hours with 1% osmium tetroxide in 0.05 M HEPES, pH 7.4. The samples were then centrifuged and washed with water 5 times, and dehydrated with acetone, using the following protocol: 15 minutes in 30%, 50%, 70%, 90% acetone each, then 100% acetone 3 changes, 30 minutes each. At the final wash step, a small amount of acetone, barely covering the pellets, was retained to avoid rehydration of the samples. The samples were then infiltrated with modified Spurr’s resin (Quetol ERL 4221 resin, Electron Microscopy Sciences, RT 14300) in a Pelco Biowave processor (Ted Pella, Inc.), as follows: 1:1 acetone:resin for 10 minutes @ 200 W – no vacuum, 1:1 acetone:resin for 5 minutes @ 200 W – vacuum 20’’ Hg (vacuum cycles with open sample container caps), 1:2 acetone:resin for 5 minutes @ 200 W – vacuum 20’’ Hg, 4 × 100% resin for 5 minutes @ 200 W – vacuum 20’’ Hg. The resin was then removed, and the sample fragments transferred to BEEM conical tip capsules prefilled with a small amount of fresh resin, resin added to fill the capsule, and capsules left to stand upright for 30 minutes to ensure that the samples sank to the bottom. The samples were then polymerized at 65 °C for 48 hours in the oven, then at left at RT for 24 hours before sectioning. 70-80 nm samples were sectioned by Leica UC / FC7 ultra-microtome (Leica Microsystems), deposited onto 300 mesh copper grids, stained with uranyl acetate / lead citrate and imaged.

### TEM immunogold sample preparation

Spores derived from the *C. difficile* R20291, and *C. difficile* Δ*cspBAC* and *C. difficile* Δ*sleC* strains were embedded in Lowicryl HM20 using HPF. 50 µL of purified spores were lightly fixed in 1 mL of 0.05% glutaraldehyde / 4% formaldehyde in 0.05 M HEPES, pH 7.4 for 30 minutes at RT. The spores were centrifuged and resuspended in 50 µL of 3% low melting point agarose. Approximately 2 µL of the spore suspension per sample was then processed with Leica EM ICE HPF apparatus (Leica Microsystems), and then freeze substituted and embedded in Lowicryl HM20 (Electron Microscopy Sciences, RT 14340), using Leica EM AFS2 freeze substitution and low temperature embedding system and Leica EM FSP freeze substitution processor. The freeze substitution protocol was adapted from Edelmann et. al. (67), with modifications, as follows: 144 hours 100% acetone at – 90 °C., temperature increase at 5 °C / hour to – 50 °C, 4 hours 100% ethanol, 4 hours 100% ethanol: 100% HM20 in 3:1 ratio, 4 hours 100% ethanol: 100% HM20 in 1:1 ratio, 4 hours 100% ethanol: 100% HM20 in 1:3 ratio, 100% HM20 2×4 hours, 100% HM20 18 hours, 100% HM20 one exchange, UV polymerization 48 hours, temperature increase 5 °C / hour to RT with UV using lamp integral to FSP, UV polymerization 48 hours at RT using dedicated UV lamp. Ultrathin sections (70-80 nm) were deposited on nickel formvar carbon coated grids, immunolabeled, stained with uranyl acetate / lead citrate, and imaged.

Spores derived from the *C. difficile* Δ*cspBAC* and *C. difficile* Δ*sleC* strains, transformed with plasmids harboring BA-FLAG and SleC-FLAG fusions were processed using PLT in Leica EM AFS2 freeze substitution and low temperature embedding system. 50 µL of purified spore samples were lightly fixed in 1 mL of 0.05% glutaraldehyde / 4% formaldehyde in 0.05 M HEPES, pH 7.4 for 30 minutes at RT. The spores were centrifuged and resuspended in 50 µL of 3 % low melting point agarose. About 10 µL of the sample were placed in a cryotubes filled with 30% ethanol prechilled to 0 °C. The protocol was as follows: 30% ethanol 1 hr, replaced with 50% ethanol, lowered the temperature to – 20 °C (5 °C / hr), 50% ethanol 1 hr, replaced with 70% ethanol, lowered the temperature to – 35 °C (5 °C / hr), 70% ethanol overnight, replaced with 90% ethanol for 1 hr, replace with 100% ethanol for 1 hr, twice, K4M 33% for 1 hr, K4M 50% for 1 hr, K4M 66% for 1 hr, K4M 100% overnight, K4M 100% 1 hr, thrice, samples placed into pre-chilled BEEM conical capsules prefilled with K4M, equilibrate at -35 °C for 1 hr, UV polymerize at -35 °C for 48 hours, UV polymerize while increasing the temperature to 24 °C at 5 °C / hr for a total of 24 hours, UV polymerize at RT for 48 hours. Ultrathin sections (70-80 nm) were deposited on nickel formvar carbon coated grids, immunolabeled, stained with uranyl acetate / lead citrate, and imaged.

### Immunolabeling

Immunolabeling of Lowicryl-embedded samples was performed using modified protocol by Dittmann et al. (68). Samples were incubated at RT, section facing down on 30 µL droplets as follows: 2 times 5 minutes 50 mM glycine in PBS, blocking solution (0.5% cold water fish skin gelatin, 0.5% bovine serum albumin, 0.01% Tween 20, in PBS) for 1 min, blocking solution for 1 hr, primary antibody diluted in blocking solution for 1 hr, 6 time blocking solution for 5 minutes, secondary antibody in blocking solution for 1 hour, 6 times blocking solution for 5 minutes, 6 times PBS for 5 minutes, 5 minutes 2% glutaraldehyde in PBS, 6 times dH_2_O for 5 minutes. For the detection of FLAG-conjugated proteins we used rabbit anti-DYKDDDDK antibody (Rockland Immunochemicals, Inc., Number 600-401-383). For the detection of CspB, CspA, CspC, and SleC we used rabbit α-CspB, α-CspA, α-CspC, and α-SleC antibodies. Secondary antibodies used were goat-anti-Rabbit IgG conjugated to 10nm gold particles (Electron Microscopy Sciences, Cat. # 25109). We also labeled the samples with secondary gold-conjugated antibody alone to test for the non-specific labeling and found virtually no labeling in each case (data not shown). For each sample immunolabeled in this way (*i*.*e*., R20291, Δ*cspBAC*, Δ*sleC*, CspBA-FLAG, and SleC-FLAG tagged samples) we imaged 100 random spores on the grid and counted the gold labels present. This was repeated at least 3 times, each time on new sections deposited on a new grid, immunolabeled in an independent trial.

### SEM sample preparation

Spores for SEM imaging of germination were prepared in the same way as the spores for TEM ultrastructure imaging, described above, up to the dehydration step. At this point the samples were processed as follows: 35 / 50 / 70 / 80% ethanol 10 min each, 1:2 hexamethyldisilazane (HMDS):ethanol for 20 minutes, 1:1 HMDS:ethanol for 20 min, 100% HMDS 2 × 20 min. At the last stage of HMDS drying, a small volume of HMDS was left in the tube, and about 10 µL of sample deposited on the circular glass cover slip. The cover slips were left to dry overnight in the fume hood in a semi-covered glass petri dish. The samples were then sputter coated with 8 nm of Pd / Pt in Cressington 208HR Sputter Coater (Cressington Scientific Instruments Ltd.) and imaged.

### TEM / SEM imaging

All ultrathin TEM sections were imaged on JEOL 1200 EX TEM (JEOL, Ltd.) at 100 kV, images were recorded on SIA-15C CCD (Scientific Instruments and Applications) camera at the resolution of 2721 × 3233 pixels using MaxImDL software (Diffraction Limited). Images were subsequently adjusted for brightness / contrast using Fiji (69).

SEM samples were imaged on Quanta 600 FEG (FEI Technologies, Inc.) equipped with a field emission gun at 20 kV, at 1024 × 943 pixels.

All equipment used is located at Texas A and M University Microscopy and Imaging Center Core Facility (RRID:SCR_022128) (70).

### Statistical analysis

Data represents results from at least 3 independent experiments, and the error bars represent standard errors of the means. One-way ANOVA followed by Tukey’s or Šídák’s multiple-comparison test, as indicated, was performed using GraphPad Prism version 9.0.2 for Windows (GraphPad Software, San Diego, California USA).

## Acknowledgments

This project was supported by awards 5R01AI116895 and 1U01AI124290 to J.A.S. from the National Institute of Allergy and Infectious Diseases. The content is solely the responsibility of the authors and does not necessarily represent the official views of the NIAID. The funders had no role in study design, data collection and interpretation, or the decision to submit the work for publication.

## References

1. Buggy BP, Wilson KH, Fekety R. 1983. Comparison of methods for recovery of Clostridium difficile from an environmental surface. Journal of Clinical Microbiology 18:348–352.

2. Jump RL, Pultz MJ, Donskey CJ. 2007. Vegetative Clostridium difficile survives in room air on moist surfaces and in gastric contents with reduced acidity: a potential mechanism to explain the association between proton pump inhibitors and C. difficile-associated diarrhea? Antimicrob Agents Chemother 51:2883–7.

3. Buffie CG, Bucci V, Stein RR, McKenney PT, Ling L, Gobourne A, No D, Liu H, Kinnebrew M, Viale A, Littmann E, van den Brink MR, Jenq RR, Taur Y, Sander C, Cross JR, Toussaint NC, Xavier JB, Pamer EG. 2015. Precision microbiome reconstitution restores bile acid mediated resistance to Clostridium difficile. Nature 517:205–8.

4. Theriot CM, Bowman AA, Young VB. 2016. Antibiotic-induced alterations of the gut microbiota alter secondary bile acid production and allow for Clostridium difficile spore germination and outgrowth in the large intestine. mSphere 1.

5. Theriot CM, Koenigsknecht MJ, Carlson PE, Jr., Hatton GE, Nelson AM, Li B, Huffnagle GB, j ZL, Young VB. 2014. Antibiotic-induced shifts in the mouse gut microbiome and metabolome increase susceptibility to Clostridium difficile infection. Nat Commun 5:3114.

6. Wilson KH, Perini F. 1988. Role of competition for nutrients in suppression of Clostridium difficile by the colonic microflora. Infect Immun 56:2610–2614.

7. Smits WK, Lyras D, Lacy DB, Wilcox MH, Kuijper EJ. 2016. Clostridium difficile infection. Nat Rev Dis Primers 2:16020.

8. Permpoonpattana P, Tolls EH, Nadem R, Tan S, Brisson A, Cutting SM. 2011. Surface layers of Clostridium difficile endospores. J Bacteriol 193:6461–70.

9. Paredes-Sabja D, Shen A, Sorg JA. 2014. Clostridium difficile spore biology: sporulation, germination, and spore structural proteins. Trends Microbiol 22:406–16.

10. Francis MB, Allen CA, Sorg JA. 2015. Spore cortex hydrolysis precedes dipicolinic acid release during Clostridium difficile spore germination. J Bacteriol 197:2276–83.

11. Setlow P. 2007. I will survive: DNA protection in bacterial spores. Trends Microbiol 15:172–80.

12. Baloh M, Sorg JA. 2021. Clostridioides difficile SpoVAD and SpoVAE interact and are required for dipicolinic acid uptake into spores. J Bacteriol 203:e0039421.

13. Jamroskovic J, Chromikova Z, List C, Bartova B, Barak I, Bernier-Latmani R. 2016. Variability in DPA and calcium content in the spores of Clostridium species. Frontiers in Microbiology 7.

14. Berendsen EM, Boekhorst J, Kuipers OP, Wells-Bennik MH. 2016. A mobile genetic element profoundly increases heat resistance of bacterial spores. ISME J 10:2633–2642.

15. Donnelly ML, Fimlaid KA, Shen A. 2016. Characterization of Clostridium difficile spores lacking either SpoVAC or dipicolinic acid synthetase. J Bacteriol 198:1694–707.

16. Francis MB, Sorg JA. 2016. Dipicolinic acid release by germinating Clostridium difficile spores occurs through a mechanosensing mechanism. mSphere 1.

17. Setlow P. 2014. Spore resistance properties. Microbiol Spectr 2.

18. Cowan AE, Koppel DE, Setlow B, Setlow P. 2003. A soluble protein is immobile in dormant spores of Bacillus subtilis but is mobile in germinated spores: implications for spore dormancy. Proc Natl Acad Sci U S A 100:4209–14.

19. Sunde EP, Setlow P, Hederstedt L, Halle B. 2009. The physical state of water in bacterial spores. Proc Natl Acad Sci U S A 106:19334–9.

20. Coullon H, Rifflet A, Wheeler R, Janoir C, Boneca IG, Candela T. 2018. N-Deacetylases required for muramic-delta-lactam production are involved in Clostridium difficile sporulation, germination, and heat resistance. J Biol Chem 293:18040–18054.

21. Diaz OR, Sayer CV, Popham DL, Shen A. 2018. Clostridium difficile lipoprotein GerS is required for cortex modification and thus spore germination. mSphere 3.

22. Makino S, Moriyama R. 2002. Hydrolysis of cortex peptidoglycan during bacterial spore germination. Med Sci Monit doi:-https://dx.doi.org/:119.

23. Paredes-Sabja D, Setlow P, Sarker MR. 2011. Germination of spores of Bacillales and Clostridiales species: mechanisms and proteins involved. Trends Microbiol 19:85–94.

24. Popham DL, Helin J, Costello CE, Setlow P. 1996. Analysis of the peptidoglycan structure of Bacillus subtilis endospores. J Bacteriol 178:6451–8.

25. Freer JH, Levinson HS. 1967. Fine structure of Bacillus megaterium during microcycle sporogenesis. J Bacteriol 94:441–57.

26. Crafts-Lighty A, Ellar DJ. 1980. The structure and function of the spore outer membrane in dormant and germinating spores of Bacillus megaterium. J Appl Bacteriol 48:135–45.

27. Driks A, Eichenberger P. 2016. The Spore Coat, The Bacterial Spore: from Molecules to Systems doi:doi:https://doi.org/10.1128/microbiolspec.TBS-0023-2016. American Society of Microbiology.

28. Paredes-Sabja D, Sarker MR. 2012. Adherence of Clostridium difficile spores to Caco-2 cells in culture. Journal of Medical Microbiology 61:1208–1218.

29. Castro-Cordova P, Mora-Uribe P, Reyes-Ramirez R, Cofre-Araneda G, Orozco-Aguilar J, Brito-Silva C, Mendoza-Leon MJ, Kuehne SA, Minton NP, Pizarro-Guajardo M, Paredes-Sabja D. 2021. Entry of spores into intestinal epithelial cells contributes to recurrence of Clostridioides difficile infection. Nat Commun 12:1140.

30. Castro-Córdova P, Mendoza-León MJ, Paredes-Sabja D. 2021. Using a ligate intestinal loop mouse model to investigate Clostridioides difficile adherence to the intestinal mucosa in aged mice. PLOS ONE 16:e0261081.

31. Romero-Rodríguez A, Troncoso-Cotal S, Guerrero-Araya E, Paredes-Sabja D, Ellermeier CD. 2020. The Clostridioides difficile cysteine-rich exosporium morphogenetic protein, CdeC, exhibits self-assembly properties that lead to organized inclusion bodies in Escherichia coli. mSphere 5:e01065–20.

32. Pizarro-Guajardo M, Calderón-Romero P, Romero-Rodríguez A, Paredes-Sabja D. 2020. Characterization of exosporium layer variability of Clostridioides difficile spores in the epidemically relevant strain R20291. Frontiers in Microbiology 11.

33. Diaz-Gonzalez F, Milano M, Olguin-Araneda V, Pizarro-Cerda J, Castro-Cordova P, Tzeng SC, Maier CS, Sarker MR, Paredes-Sabja D. 2015. Protein composition of the outermost exosporium-like layer of Clostridium difficile 630 spores. J Proteomics 123:1–13.

34. Shen A. 2020. Clostridioides difficile spore formation and germination: new insights and opportunities for intervention. Annual Review of Microbiology 74:545–566.

35. Sorg JA, Sonenshein AL. 2008. Bile salts and glycine as cogerminants for Clostridium difficile spores. J Bacteriol 190:2505–12.

36. Sorg JA, Sonenshein AL. 2009. Chenodeoxycholate is an inhibitor of Clostridium difficile spore germination. J Bacteriol 191:1115–7.

37. Ridlon JM, Kang DJ, Hylemon PB. 2006. Bile salt biotransformations by human intestinal bacteria. J Lipid Res 47:241–59.

38. Ridlon JM, Kang DJ, Hylemon PB, Bajaj JS. 2014. Bile acids and the gut microbiome. Curr Opin Gastroenterol 30:332–8.

39. Francis MB, Allen CA, Sorg JA. 2013. Muricholic acids inhibit Clostridium difficile spore germination and growth. PLoS One 8:e73653.

40. Ramirez N, Liggins M, Abel-Santos E. 2010. Kinetic evidence for the presence of putative germination receptors in Clostridium difficile spores. J Bacteriol 192:4215–22.

41. Howerton A, Ramirez N, Abel-Santos E. 2011. Mapping interactions between germinants and Clostridium difficile spores. J Bacteriol 193:274–82.

42. Bhattacharjee D, Francis MB, Ding X, McAllister KN, Shrestha R, Sorg JA. 2016. Reexamining the germination phenotypes of several Clostridium difficile strains suggests another role for the CspC germinant receptor. Journal of Bacteriology 198:777–786.

43. Shrestha R, Sorg JA. 2018. Hierarchical recognition of amino acid co-germinants during Clostridioides difficile spore germination. Anaerobe 49:41–47.

44. Kochan TJ, Somers MJ, Kaiser AM, Shoshiev MS, Hagan AK, Hastie JL, Giordano NP, Smith AD, Schubert AM, Carlson PE, Jr., Hanna PC. 2017. Intestinal calcium and bile salts facilitate germination of Clostridium difficile spores. PLoS Pathog 13:e1006443.

45. Leslie JL, Jenior ML, Vendrov KC, Standke AK, Barron MR, O’Brien TJ, Unverdorben L, Thaprawat P, Bergin IL, Schloss PD, Young VB. 2021. Protection from lethal Clostridioides difficile infection via intraspecies competition for cogerminant. mBio 12.

46. Setlow P, Wang S, Li YQ. 2017. Germination of spores of the orders Bacillales and Clostridiales. Annu Rev Microbiol 71:459–477.

47. Donnelly ML, Forster ER, Rohlfing AE, Shen A. 2020. Differential effects of ‘resurrecting’ Csp pseudoproteases during Clostridioides difficile spore germination. Biochemical Journal 477:1459–1478.

48. Rohlfing AE, Eckenroth BE, Forster ER, Kevorkian Y, Donnelly ML, Benito de la Puebla H, Doublie S, Shen A. 2019. The CspC pseudoprotease regulates germination of Clostridioides difficile spores in response to multiple environmental signals. PLoS Genet 15:e1008224.

49. Kevorkian Y, Shen A. 2017. Revisiting the role of Csp family proteins in regulating Clostridium difficile spore germination. J Bacteriol 199.

50. Kevorkian Y, Shirley DJ, Shen A. 2016. Regulation of Clostridium difficile spore germination by the CspA pseudoprotease domain. Biochimie 122:243–54.

51. Adams CM, Eckenroth BE, Putnam EE, Doublie S, Shen A. 2013. Structural and functional analysis of the CspB protease required for Clostridium spore germination. PLoS Pathog 9:e1003165.

52. Shimamoto S, Moriyama R, Sugimoto K, Miyata S, Makino S. 2001. Partial characterization of an enzyme fraction with protease activity which converts the spore peptidoglycan hydrolase (SleC) precursor to an active enzyme during germination of Clostridium perfringens S40 spores and analysis of a gene cluster involved in the activity. J Bacteriol 183:3742–51.

53. Paredes-Sabja D, Setlow P, Sarker MR. 2009. The protease CspB is essential for initiation of cortex hydrolysis and dipicolinic acid (DPA) release during germination of spores of Clostridium perfringens type A food poisoning isolates. Microbiology 155:3464–72.

54. Shrestha R, Cochran AM, Sorg JA. 2019. The requirement for co-germinants during Clostridium difficile spore germination is influenced by mutations in yabG and cspA. PLoS Pathog 15:e1007681.

55. Sorg JA, Sonenshein AL. 2010. Inhibiting the initiation of Clostridium difficile spore germination using analogs of chenodeoxycholic acid, a bile acid. J Bacteriol 192:4983–90.

56. Kochan TJ, Foley MH, Shoshiev MS, Somers MJ, Carlson PE, Hanna PC. 2018. Updates to Clostridium difficile spore germination. J Bacteriol 200.

57. Fimlaid KA, Jensen O, Donnelly ML, Francis MB, Sorg JA, Shen A. 2015. Identification of a novel lipoprotein regulator of Clostridium difficile spore germination. PLoS Pathog 11:e1005239.

58. Francis MB, Sorg JA. 2016. Detecting cortex fragments during bacterial spore germination. J Vis Exp doi:10.3791/54146.

59. Heeg D, Burns DA, Cartman ST, Minton NP. 2012. Spores of Clostridium difficile clinical isolates display a diverse germination response to bile salts. PLoS One 7:e32381.

60. Wang S, Shen A, Setlow P, Li Y-q, Boer Pd. 2015. Characterization of the dynamic germination of individual Clostridium difficile spores using Raman spectroscopy and differential interference contrast microscopy. Journal of Bacteriology 197:2361–2373.

61. Miyata S, Moriyama R, Miyahara N, Makino S. 1995. A gene (sleC) encoding a spore-cortex-lytic enzyme from Clostridium perfringens S40 spores; cloning, sequence analysis and molecular characterization. Microbiology 141 (Pt 10):2643–50.

62. Miyata S, Kozuka S, Yasuda Y, Chen Y, Moriyama R, Tochikubo K, Makino S. 1997. Localization of germination-specific spore-lytic enzymes in Clostridium perfringens S40 spores detected by immunoelectron microscopy. FEMS Microbiol Lett 152:243–7.

63. Banawas S, Korza G, Paredes-Sabja D, Li Y, Hao B, Setlow P, Sarker MR. 2015. Location and stoichiometry of the protease CspB and the cortex-lytic enzyme SleC in Clostridium perfringens spores. Food Microbiol 50:83–7.

64. Francis MB, Allen CA, Shrestha R, Sorg JA. 2013. Bile acid recognition by the Clostridium difficile germinant receptor, CspC, is important for establishing infection. PLoS Pathog 9:e1003356.

65. Hanahan D. 1983. Studies on transformation of Escherichia coli with plasmids. J Mol Biol 166:557–80.

66. Gibson DG, Young L, Chuang RY, Venter JC, Hutchison CA, 3rd, Smith HO. 2009. Enzymatic assembly of DNA molecules up to several hundred kilobases. Nat Methods 6:343–5.

67. Edelmann L. 1991. Freeze-substitution and the preservation of diffusible ions. Journal of Microscopy 161:217–228.

68. Dittmann C, Han H-M, Grabenbauer M, Laue M. 2015. Dormant Bacillus spores protect their DNA in crystalline nucleoids against environmental stress. Journal of Structural Biology 191:156–164.

69. Schindelin J, Arganda-Carreras I, Frise E, Kaynig V, Longair M, Pietzsch T, Preibisch S, Rueden C, Saalfeld S, Schmid B, Tinevez J-Y, White DJ, Hartenstein V, Eliceiri K, Tomancak P, Cardona A. 2012. Fiji: an open-source platform for biological-image analysis. Nature Methods 9:676–682.

70. Resource. Texas A and M University Microscopy and Imaging Center Core Facility (RRID:SCR_022128).

71. Ma NJ, Moonan DW, Isaacs FJ. 2014. Precise manipulation of bacterial chromosomes by conjugative assembly genome engineering. Nature Protocols 9:2285–2300.

72. Heap JT, Pennington OJ, Cartman ST, Minton NP. 2009. A modular system for Clostridium shuttle plasmids. J Microbiol Methods 78:79–85.

73. McAllister KN, Bouillaut L, Kahn JN, Self WT, Sorg JA. 2017. Using CRISPR-Cas9-mediated genome editing to generate C. difficile mutants defective in selenoproteins synthesis. Scientific Reports 7:14672.

74. Kaus GM, Snyder LF, Müh U, Flores MJ, Popham DL, Ellermeier CD. 2020. Lysozyme resistance in Clostridioides difficile is dependent on two peptidoglycan deacetylases. Journal of bacteriology 202:e00421–20.

